# Deciphering the code of viral-host adaptation through maximum entropy models

**DOI:** 10.1101/2023.10.28.564530

**Authors:** Andrea Di Gioacchino, Benjamin D. Greenbaum, Remi Monasson, Simona Cocco

## Abstract

Understanding how the genome of a virus evolves depending on the host it infects is an important question that challenges our knowledge about several mechanisms of host-pathogen interactions, including mutational signatures, innate immunity, and codon optimization. A key facet of this general topic is the study of viral genome evolution after a host-jumping event, a topic which has experienced a surge in interest due to the fight against emerging pathogens such as SARS-CoV-2. In this work, we tackle this question by introducing a new method to learn Maximum Entropy Nucleotide Bias models (MENB) reflecting single, di- and tri-nucleotide usage, which can be trained from viral sequences that infect a given host. We show that both the viral family and the host leave a fingerprint in nucleotide usages which MENB models decode. When the task is to classify both the host and the viral family for a sequence of unknown viral origin MENB models outperform state of the art methods based on deep neural networks. We further demonstrate the generative properties of the proposed framework, presenting an example where we change the nucleotide composition of the 1918 H1N1 Influenza A sequence without changing its protein sequence, while manipulating the nucleotide usage, by diminishing its CpG content. Finally we consider two well-known cases of zoonotic jumps, for the H1N1 Influenza A and for the SARS-CoV-2 viruses, and show that our method can be used to track the adaptation to the new host and to shed light on the more relevant selective pressures which have acted on motif usage during this process. Our work has wide-ranging applications, including integration into metagenomic studies to identify hosts for diverse viruses, surveillance of emerging pathogens, prediction of synonymous mutations that effect immunogenicity during viral evolution in a new host, and the estimation of putative evolutionary ages for viral sequences in similar scenarios. Additionally, the computational frame-work introduced here can be used to assist vaccine design by tuning motif usage with fine-grained control.

**Author summary:** In our research, we delved into the fascinating world of viruses and their genetic changes when they jump from one host to another, a critical topic in the study of emerging pathogens. We developed a novel computational method to capture how viruses change the nucleotide usage of their genes when they infect different hosts. We found that viruses from various families have unique strategies for tuning their nucleotide usage when they infect the same host. Our model could accurately pinpoint which host a viral sequence came from, even when the sequence was vastly different from the ones we trained on. We demonstrated the power of our method by altering the nucleotide usage of an RNA sequence without affecting the protein it encodes, providing a proof-of-concept of a method that can be used to design better RNA vaccines or to fine-tune other nucleic acid-based therapies. Moreover the framework we introduce can help tracking emerging pathogens, predicting synonymous mutations in the adaptation to a new host and estimating how long viral sequences have been evolving in it. Overall, our work sheds light on the intricate interactions between viruses and their hosts.

## 1. Introduction

The recent COVID-19 pandemic inspired the scientific community to investigate zoonotic transmission of viruses [Parrish et al., 2008, Andersen et al., 2020] and the subsequent evolutionary dynamics of viral adaptation to a new host. Several experimental [Starr et al., 2020, Moulana et al., 2022] and computational [Rodriguez-Rivas et al., 2022, Tubiana et al., 2022] investigations pointed out the impact of amino-acid mutations in the spike glycoprotein and their effects on its interaction with the human ACE2 receptor, which conferred a fitness advantage and resulted in selective sweeps of new variants [Kang et al., 2021, Lee et al., 2022].

Another fundamental question is identifying Pathogen-Associated Molecular Patterns (PAMPs) in a viral sequence [Akira and Hemmi, 2003] and predicting how the virus changed those patterns to adapt to the human environment and to alter innate immune recognition and response. This topic had been previously explored for the H1N1 strain of the 1918 H1N1 influenza pandemic. In this context it has been shown that the viral genome evolved in a predictable way to lose CpG motifs (a cytosine followed by a guanine in the 5’-to-3’ sense) after entering its human host from an avian reservoir [Greenbaum et al., 2008, Greenbaum et al., 2014]. This observation, together with the fact that most human-infecting viruses have a low abundance of CpG motifs, was followed by the identification of the CpG-dependent receptor specificity of the human Zinc-finger Antiviral Protein (ZAP, coded by ZC3HAV1 gene) [Gao et al., 2002, Takata et al., 2017], implying such approaches can identify recognition sites by host anti-viral restriction factors. Similar analyses for the early evolution of SARS-CoV-have been carried out [Di Gioacchino et al., 2021, Kumar et al., 2022], showing a similar pressure to reduce CpG motifs in CpG-rich regions of the viral genome. Finally, understanding and controlling the impact of a foreign RNA sequence on the stimulation of the innate immune response has an important application in DNA and RNA vaccine design in order to avoid over-stimulating the host innate reaction to nucleic acids in the vaccine[Zhang et al., 2023], while also optimizing for features such as codon bias [Pardi et al., 2018].

These questions are facets of the fundamental problem of determining how the interaction of a virus with its host is imprinted upon evolving viral genomes. This topic has been considered in several contexts [Hall et al., 2013, Bloom et al., 2023], demonstrating that viruses of the same family accumulate mutations to use similar nucleotide patterns when they evolve in interaction with a specific host. This idea has been in turn the cornerstone of a fruitful series of works aimed at determining the host of a virus from its genome. Remarkably, it has been shown that methods that do not resort to sequence alignment perform, for this specific task, comparably well with alignment-based methods [Li and Sun, 2018]. These methods typically rely on using machine learning based on the frequencies of *k*-mers (subsequences of length *k*) up to a given length *k*_max_, either alone [Tang et al., 2015, Brierley and Fowler, 2021], together with other features such as physical-chemical properties of amino-acids [Young et al., 2020], or using a hybrid method that integrates alignment-based features [Babayan et al., 2018]. Recently, techniques based on deep neural networks have been suggested to solve the task of finding the correct host of a given virus, completely by-passing the choice of the features used for a model [Mock et al., 2020]. While most of these methods can give remarkable classification performances, there is a pressing need for techniques that are effective at the classification task while remaining at the same time simple to use and interpretable. The latter point is particularly important to increase our molecular understanding of the evolutionary processes that a virus undergoes after an host switch, which can then be targeted by a antiviral therapies during a zoonitic transmission.

In this work, we address all these issues by taking a novel approach: we build a maximum entropy model whose parameters are inferred to capture short-range (up to 3-mers) nucleotide usage patterns in viral genome sequences. Maximum-entropy models have been already used in several contexts, such as for protein sequences [Morcos et al., 2011, Cocco et al., 2018,Mayer et al., 2022], neuronal spiking activity [Tavoni et al., 2017, Ferrari et al., 2017] and social dynamics [Bialek et al., 2012, Chen et al., 2022], demonstrating the effectiveness and flexibility of this approach. In the context of viral evolution and identification of PAMPS in RNA sequences the approach introduced here extends the selective force model previously introduced [Greenbaum et al., 2014, Tanne et al., 2015, Di Gioacchino et al., 2021] which reproduced the motif usage of a particular k-mers only: CpG dinucleotide and other individual motifs. In analogy to *k*-mer based methods our model does not require any alignment or annotation of the genetic sequence under analysis. We show our technique is simple but extremely effective to tackle the host classification task, resulting in performances comparable with deep neural network models or, in the more challenging setting where no phylogenetic information is available, superior in its discrimination capability.

## 2. Results

### 2.1. MENB: a model for host and viral origin classification

Our unsupervised learning model, MENB, infers parameters associated for each *k*-mer up to *k* = 3 and defines a probability distribution on viral sequences (of a fixed length), in such a way that the expected *k*-mer frequencies from this distribution match with those observed in the training data. As shown in Methods Sec. 5.1.1, this results in the following probability distribution for a viral sequence ***s***:

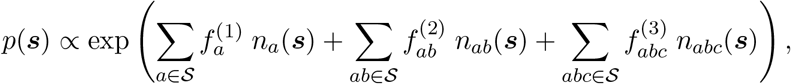

where 𝒮 is the set of nucleotides, *n*_m_(***s***) is the number of times the motif m is present in ***s***, and the parameters indicated by *f* are the “forces” [Greenbaum et al., 2014] to be inferred from the training data.

To train our model we collected viral sequences from the BV-BRC database [Olson et al., 2022], and filtered the data for sequences of three host classes: human, avian and swine viruses. We required at least 150 (different) viral genomes for each host class, and this left us with 4 viral families: *Coronaviridae, Flaviviridae, Picornaviridae*, and *Orthomyxoviridae*(focusing on Influenza A alone). We stress that such number of sequences is in principle not necessary to train our models: a single sequence (of sufficient length) is enough, provided that the number of motifs observed in that sequence is representative. To avoid biases in choosing this reference sequence, however, we decided to train the models on sets of 100 sequences (the remaining sequences are used as test set). We then test the model in the task of host classification from a viral sequence. We consider three strategies to assign an host to a given viral sequence. In the simplest one, called “MENB-H”, for each host *h* we grouped together the sequences belonging to different viral families and trained a single MENB model that approximates the probability *p*(*s*|*h*). Given a new sequence *s*, we can therefore estimate the probability of it coming from host *h* using Bayes formula *p*(*h*|*s*) ∝ *p*(*s*|*h*) *p*(*h*), where *p*(*h*) is a prior that we will consider uniform over the host distribution.

To introduce a more complex strategy we start by training a set of MENB models *p*(*s*|*h, v*) at fixed viral family *v* and host *h*. As in the previous case, we can then obtain the probability of a sequence to be associated to a host-virus, (*h, v*), pair as *p*(*h, v*|*s*) ∝ *p*(*s*|*h, v*) *p*(*h, v*). If we know the viral origin (*v*_0_) of the test sequence we can limit ourselves to compare models trained for that family on different host, a strategy that we name “MENB-H|V”, and by assuming an uniform prior *p*(*h, v*_0_) we obtain *p*(*h*|*s, v*_0_) ∝ *p*(*s*|*h, v*_0_). If, on the contrary, we ignore the viral family of the sequence we can then sum over the different viral families to have a probability a virus is associated with a given host, a strategy that we will call “MENB-H,V”. By assuming again a uniform prior we obtain *p*(*h*|*s*) ∝ ∑_*v*_ *p*(*s*|*h, v*). Remarkably, for all viral genomes analyzed in this work, there is a unique term that contributes much more than all the others to the above summation. Hence we can associate to a viral sequence a specific host as the most likely origin, and likewise guess the viral family from the term that mostly contributes to the probability of that host.

The results of the host classification task on test viral sequences, after having trained the models using the three strategies (“MENB-H”, “MENB-H,V”, “MENB-H|V”) discussed are displayed in in Fig. 1A. We first notice that the viral agnostic models, MENB-H, has a low accuracy: the accuracy averaged over the viral families is about 51%, blue dashed line), only marginally better than random guessing (33%, black dashed line), with performances comparable to random guessing for *Coronaviridae* and *Orthomyxoviridae*. Similar results have been observed elsewhere [Mock et al., 2020].

**Figure 1.**
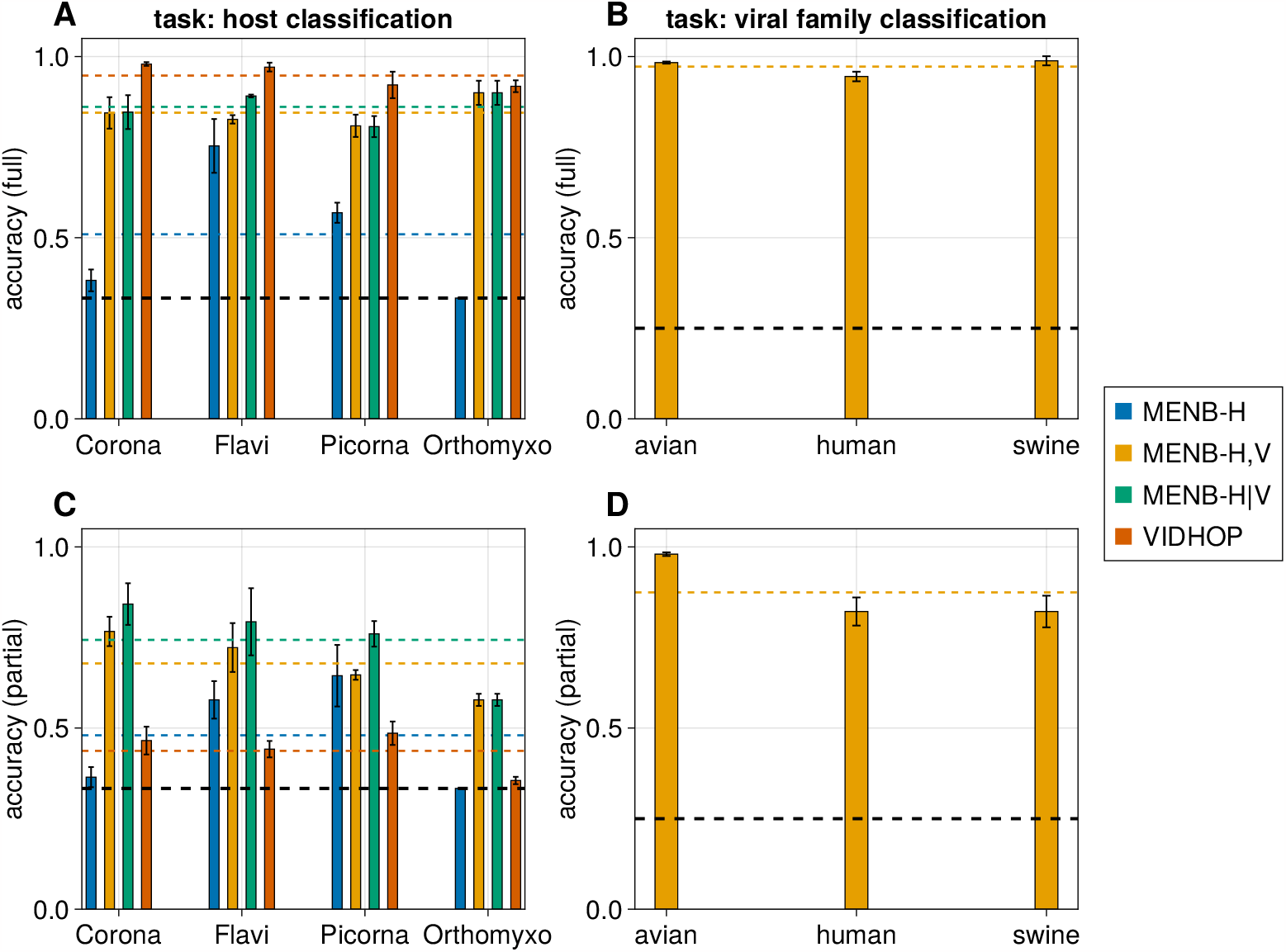
MENB models can predict host and viral family of viral genomes. A: accuracy of MENB models trained on all viruses with the same host (blue bars) and on all virus-host pairs (orange bars) on the host classification task on the test set of viral genomes; green bars are obtained using the same models used for the orange bar, but using only the correct viral family of the target viral genome, and red bars are the accuracy of the host classification task in this same setting with the VIDHOP algorithm. B: accuracy of MENB models trained on all virus-host pairs in determining the correct viral family for the target test genome as the one that mostly contributes in the host classification. C: same as A, but the training is done on the first half of the genome (for *Coronaviridae, Flaviviridae, Picornaviridae*) or on all segments but PB2 (for *Orthomyxoviridae*), and the test is done on the remaining part of the sequence. D: same as B, with the same task as described in C.

A possible explanation for the failure of this viral-agnostic host inference strategy is that viral genomes are highly constrained (for instance, they need to code for multiple, sometimes overlapping protein sequences while interacting with viral proteins for encapsulation), hence not free to evolve to change their nucleotide usage in a way that depends uniquely upon the host. Such explanation is confirmed by the improved performances obtained when learning viral-families dependent models for each hosts (“MENB-V,H”), and marginalizing over viral families to find the most probable host. “MENB-V,H” gives an average performance in classification of (85%, orange bars in Fig. 1A). Moreover when comparing (Fig. 1B) the values of *v* that give the largest contribution to the sum with the real viral families. We find an average accuracy of about 97%, confirming that the “MENB-V,H” strategy is able to predict, with a very good accuracy, both the host and the viral family of a new sequence.

### 2.2. Comparison of MENB with other approaches

Given the performance of MENB models for the host classification task, a natural question is how it compares with other state-of-the-art approaches. To answer this, we considered VIDHOP [Mock et al., 2020], a deep-neural network designed specifically for this task which can be obtained from a public code repository re-trained by any user. The authors in [Mock et al., 2020] noticed that their algorithm could not generalize to viruses of different families, so they designed VIDHOP to work at fixed viral family. As we demonstrated, MENB can in principle work without information about the viral family of the target sequence, but to make the comparison fairer we modified our approach to use MENB models to assess the host of viral sequences at fixed viral family: we considered as hosts directly the arg max_*h*_ *p*(*h, v*|*s*), where the correct viral family *v* is used instead of summing on all possible families. We retrained VIDHOP and MENB on the same sequences, and compared their performances. As expected from the higher complexity (in terms of number of learnable parameters) of VIDHOP, its performances are better than MENB in most cases and in particular for *Coronaviridae*, while being very similar for *Orthomyxoviridae*, as shown in Fig. 1A (green and red bars). On the other hand, VIDHOP requires many more resources (in terms of time and computational power) with respect to MENB (for instance, for each viral family VIDHOP requires about 1 hour on a 56-core CPU, while MENB requires less than 5 minutes on 3 cores).

We then wanted to confirm that the host classification results we obtained with MENB models are actually related to viral adaptation to their hosts, and not caused by spurious effects such as phylogenetic correlations that lead to strong similarity of sequences in the training and test set. We therefore designed a more difficult classification task based on out-of-distribution data points: we trained our model on a part of the viral se-quences (the first half for *Coronaviridae, Flaviviridae* and *Picornaviridae*, and on all segments but PB2 for *Orthomyxoviridae*), and used it to determine the host from the other part of the sequences. In this way the classification is performed on sequences that are completely different (in terms of edit distance) from those used during training, but as shown in Fig. 1C and D, the model can still determine quite precisely the viral family of the test part of the sequences (the average accuracy is about 89%), and performs much better than a random classifier in determining the host (the average accuracy is about 67%), although the performances are degraded with respect to those obtained with full sequences. Remarkably, in this test MENB performs sensibly better than VIDHOP, whose results are only marginally superior than those of a random classifier (black dashed line in the plot). It is therefore reasonable to expect that the extremely good performance of VIDHOP on full sequences relies on the large similarity between training and test sequences, even if cross-validation during training is used to select the best model on a validation dataset.

In general, the performance of MENB models derive from the differences between the probability distributions over viral sequences that each model learns. In Fig. 2 we show the symmetrized Kullback-Leibler (KL) divergence (for a definition, see Methods Sec. 5.1.3) between each pair of distributions. Remarkably, models trained on viruses infecting the same host encode far more different probability distributions than models trained on viruses of the same family, suggesting that the nucleotide usage is more driven by phylogenetic correlations than by host adaptation. This is compatible with the much greater performances of the MENB models in discriminating viral families rather than hosts, and the smaller divergences within viral families ultimately justify the choice of using the “MENB-H,V” strategy. Moreover, we notice that *Orthomyxoviridae* viruses have smaller differences between hosts with respect to other viral families, probably because of their tendency to commonly undergo reassortments with segments of viruses adapted to different hosts.

**Figure 2.**
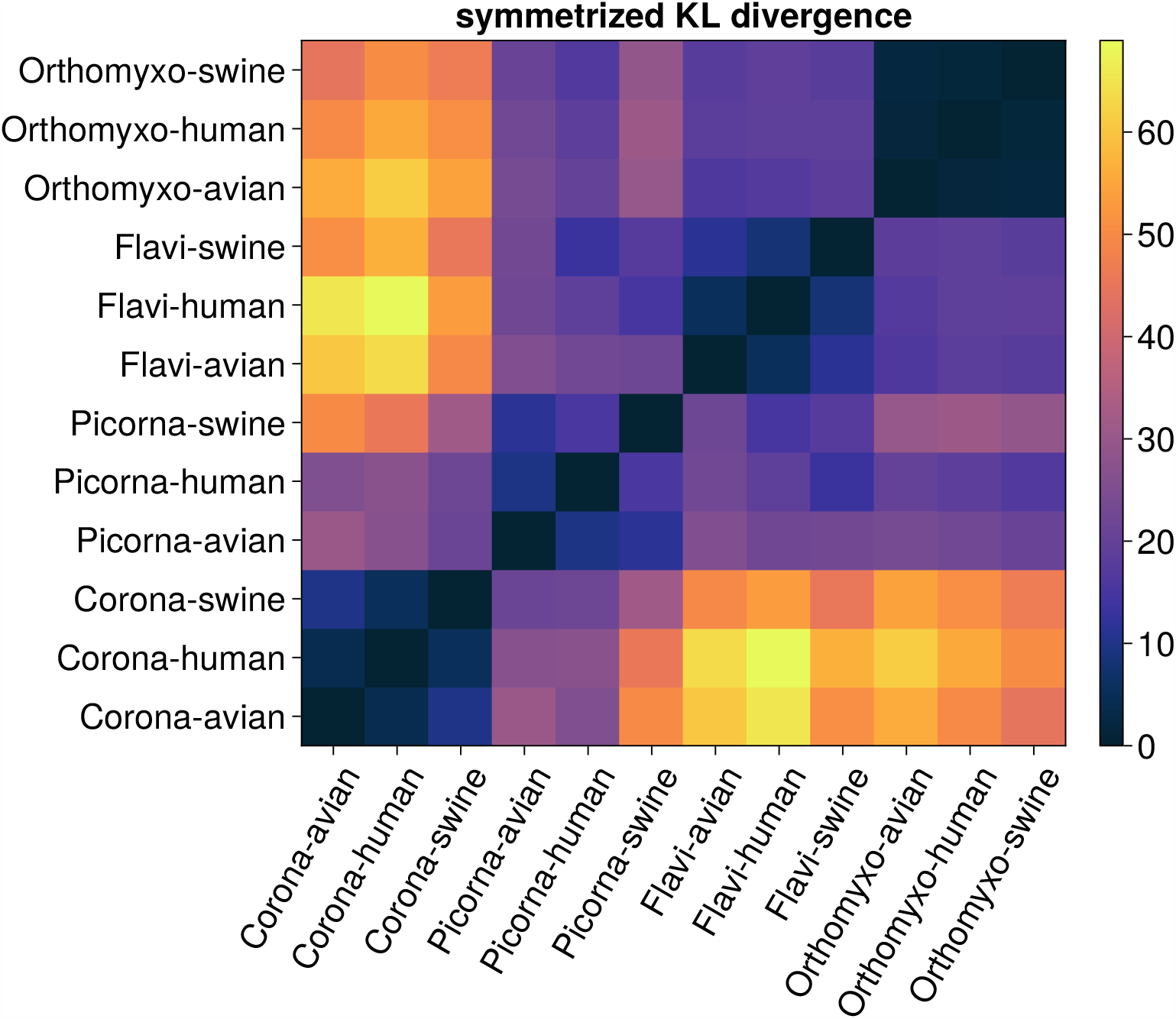
Viruses infecting the same host use nucleotides in different ways. Symmetrized Kullback-Leibler divergences between all (full) MENB model pairs considered in this work. The divergence is computed with respect to sequences having an arbitrary length of 1000 nucleotides, see Methods Sec. 5.1.3.

### 2.3. Generative power of MENB models

In Fig. 3 we focus on the human *Orthomyxoviridae* viral sequences and we show that the MENB model reproduces, as expected, the 1-, 2- and 3-mer statistics of the training set (Suppl. Fig. 5). Moreover, it generalizes to new sequences in the test set, which are not used for the training, when full genomes are used (Fig. 3A). These performances are only slightly degraded when a fraction of each genome is used in the training test and the test set contains new sequences and the unseen part of the genome (Fig. 3C), further showing how nucleotide usage biases encompass the full viral sequences and can be learned from a fraction of them.

**Figure 3.**
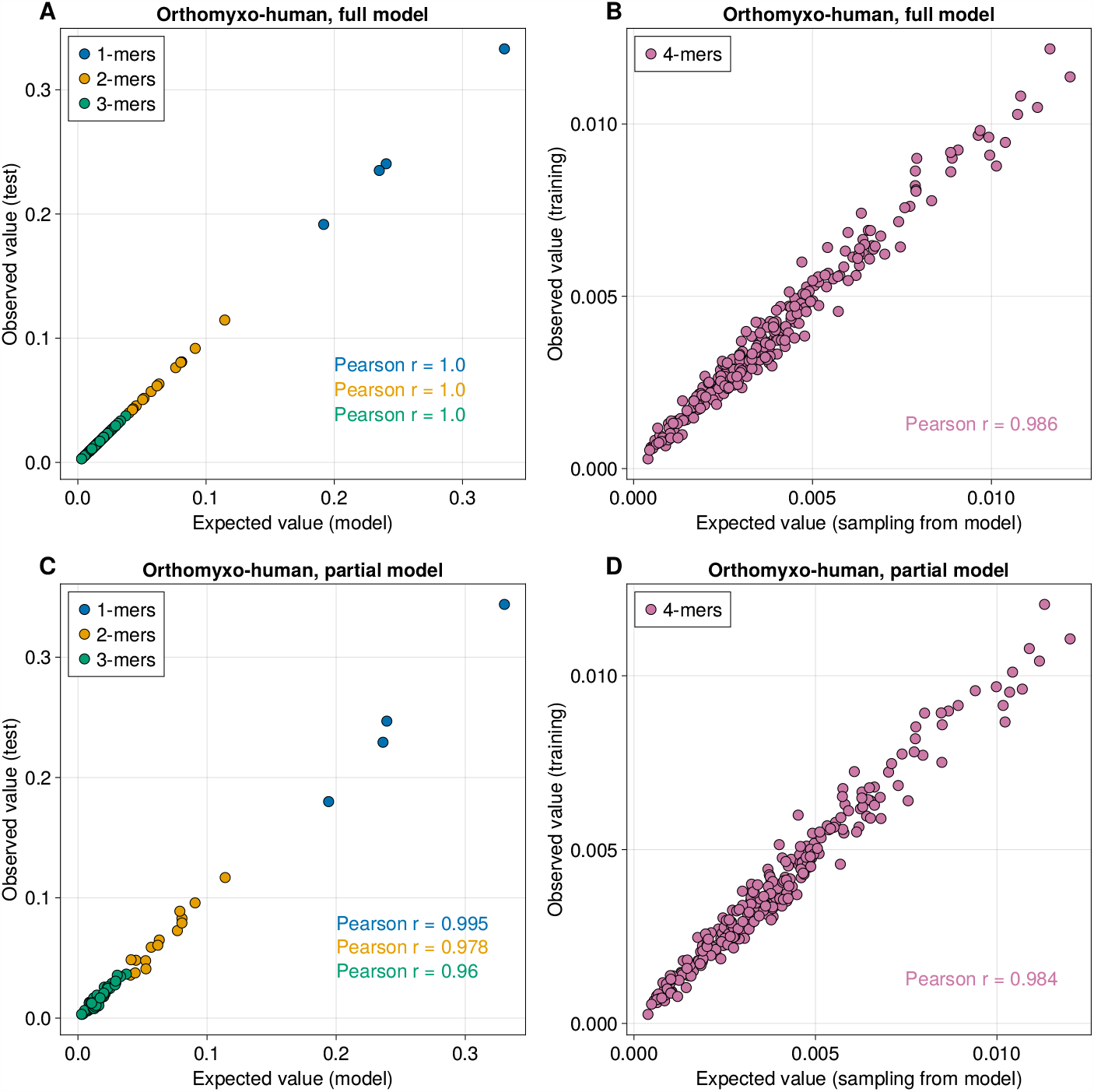
MENB models generalize well to test sequences and higher-order motifs. A: Frequency of nucleotides, 2-mers and 3-mers observed in the test set of full human *Orthomyxoviridae* sequences versus the value obtained analytically from the inferred MENB model. B: Same as A for the MENB model trained on human *Orthomyxoviridae* sequences without the segment coding for PB2. C: Frequency of 4-mers observed in the training set of full human *Orthomyxoviridae* sequences versus the value obtained from sequenced sampled from the inferred MENB model. D: Same as C for the MENB model trained on human *Orthomyxoviridae* sequences without the segment coding for PB2.

The MENB models are trained to reproduce the frequency of 1-, 2-, and 3-mers observed in the training dataset; we next investigated how well these models reproduce higher order statistics. To do so, we sampled sequences from the probability distribution encoded by MENB models (using a standard Metropolis–Hastings algorithm) and compared to the 4-mer frequencies observed in these sampled sequences with those of the training dataset. In Fig. 3B,D we show that MENB model almost perfectly capture the 4-mer statistics.

In Fig. 4 we further show how we can leverage MENB models together with the Metropolis–Hastings sampling algorithm to change the nucleotide usage of a protein-coding sequence, while keeping fixed its amino-acid sequence. As an illustration, we considered the PB2 coding region of the 1918 H1N1 strain and wanted to reduce its number of PAMP associated CpG motifs [Greenbaum et al., 2008, Greenbaum et al., 2014].

**Figure 4.**
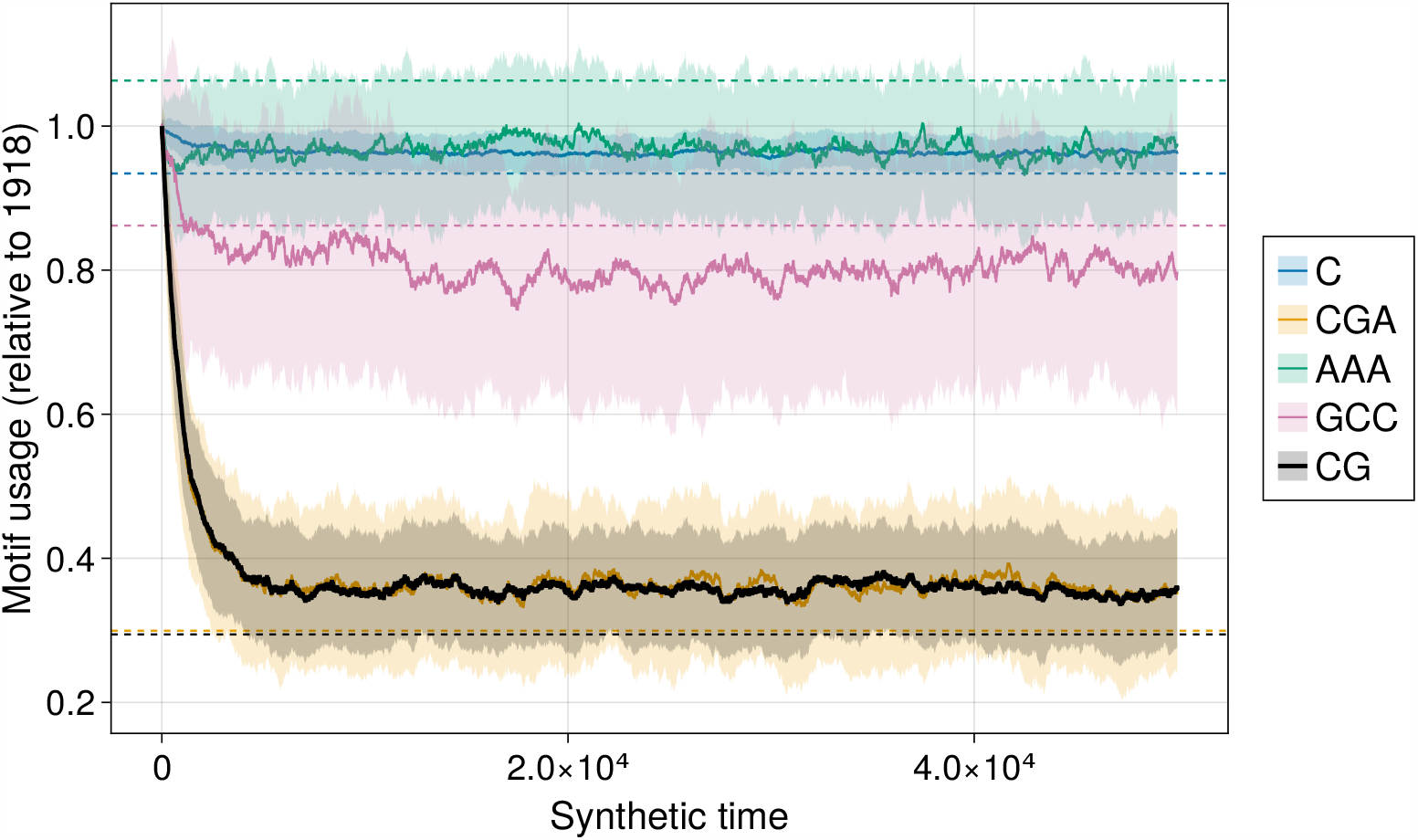
MENB models can be used to design new sequences coding for the same proteins and with different nucleotide usage. Motif usage (relative to the original sequence) during synthetic evolution of the PB2 coding sequence from 1918 H1N1 strain under a MENB model that enforces a lower usage of CpG dinucleotides. Solid lines are obtained as averages of 100 independent evolutions, and shaded areas denote one standard deviation. Dashed lines denote the expected motifs usage without the constrain to code for the PB2 protein of the 1918 H1N1 strain.

We thus synthetically evolved the 1918 sequence by the Metropolis–Hasting dynamics and using the MENB the force parameters inferred from the 1918 sequence apart from *f*_*CpG*_ which we fixed to *f*_*CpG*_ = −1.9. Such *f*_*CpG*_ value is close to the average value in the human genome and is sensibly lower than the one in the original H1N1 strain (*f*_*CpG*_ = −0.6). The original amino-acid content of the 1918 sequence was kept by accepting only synonymous mutations in the Metropolis–Hasting sampling dynamics [Chatenay et al., 2017]. As shown in Fig. 4 this resulted in a global change of the nucleotide content of the sequence, where CpG dinucleotides and CpG-containing 3-mers are mostly affected, while other dimers and trimers are generally conserved, see Suppl. Fig. 8. Generation of synthetic sequences under fixed constraint on other motifs can be analogously carried on by changing the corresponding forces.

### 2.4. Viruses adapt to their host after hosts jumps: Applications to H1N1 influenza and SARS-CoV-2

We demonstrated that our model can infer the host of viral sequences from their nucleotide statistics alone. Here we show the model describes the evolution of a viral strain after a host jump. We start with the case of 1918 H1N1 influenza pandemics: we collected all PB2 segments available in our dataset associated to the H1N1 strain up to 2008. It is commonly accepted that the pandemics originated with a jump from avian to human hosts [Taubenberger et al., 2005]. To compare the two hosts we will use in our analysis the human and the avian model trained, for each host, on all the segments of influenza viruses excluding PB2 to avoid potential overfitting. Before assigning sequences to their host, we built a phylogenetic tree, on a random subsample of up to 20 sequences per year, using Nextstrain [Hadfield et al., 2018]. Fig. 5A shows the log-probability difference between the influenza-human and influenza-avian MEMB models at fixed viral family as a function of time since 1918. The log-probability difference allows classification of the host, similarly to the host classification task with MENB-H,V in Fig.1 from the sequences sampled over time but also from the reconstructed roots along the phylogenetic tree. We observe that the maximum-entropy model is misled in the assessment of the host of the 1918 PB2 segments (left side of Fig. 5A), which is wrongly classified as an avian virus, while being sampled in humans. This mislcassification is a clear signature of the host jump which had just occurred and originated the 1918 pandemic. The classification changes with time: as the virus evolve in contact with the human host, the model assigns to it higher log-probability differences, giving equal scores to human and avian origin around 1950. For more recent samples the model is more and more confident about the human classification. Quite remarkably, the log-probability score introduced here works as a sort of “molecular clock”, by steadily increasing as the virus adapts to the new host. Similar results are obtained also by a simple model only reflecting the nucleotide usage or also including the CpG forces [Greenbaum et al., 2014] (Suppl. Fig. 4), although in these cases the difference of log-probability between the two models is less pronounced, confirming that host adaptation takes place at different order on motif’s usage.

**Figure 5.**
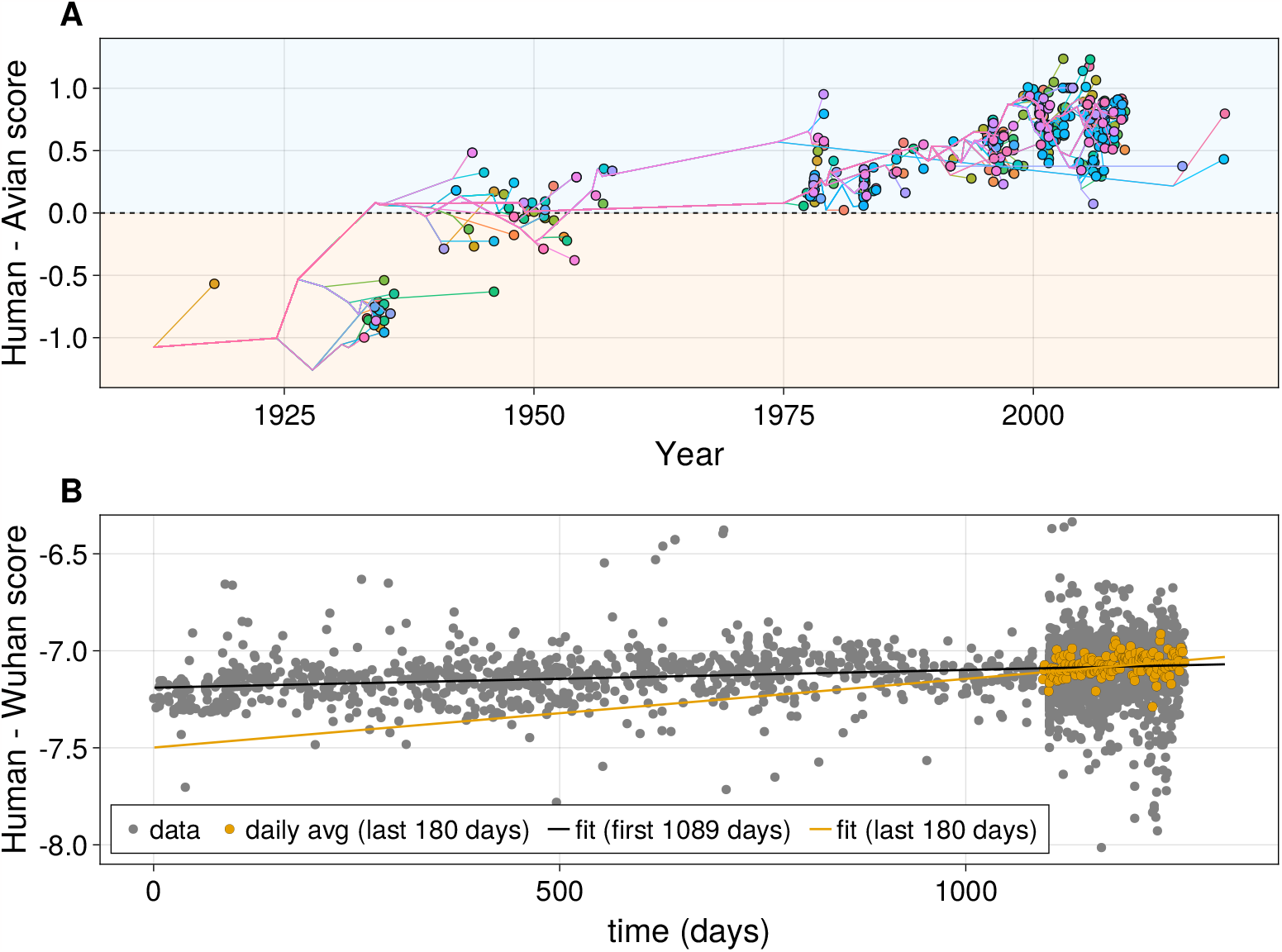
MENB models can be used to quantify host adaptation dynamics after host jumps. A: Scatter plot of loglikelihood differences of the MENB *Orthomyxoviridae* human and avian models versus time of H1N1 Influenza A sequences. The colored lines are the reconstructed paths of the inferred phylogenetic tree that connect the root to each leaf (observed sequence), and the score versus inferred time is plotted also for the internal node (inferred) sequences. B: Scatter plot of loglikelihood differences of the MENB *Coronaviridae* model versus a MENB model trained on the original Wuhan SARS-CoV-2 sequence versus time from December 26th 2019. The black line is a linear fit on the first 1089 days (slope: 9 10^*−*5^, p-value: 10^*−*9^), the orange line is a linear fit of the last 180 days (slope: 3.5 10^*−*4^, p-value: 10^*−*7^). To ease the visualization of the increasing trend of the score difference in the last 180 days, daily averages of the score differences are plotted as orange points.

As a final application of our MENB models, we turned to the SARS-CoV-2 virus. We wanted to check if we can see hints of host adaptation as for the 1918 H1N1 virus. This case is different from H1N1 as the original host of SARS-CoV-2 is currently unknown and subject of scientific debate [Andersen et al., 2020]; we have therefore assumed that the original Wuhan sequence is representative of the (unknown) previous host and build its MEMB model from this unique sequence, while building the model for SARS-Cov-2 in human host from the sequences collected during the recent pandemic waves and collected in Nextstrain [Hadfield et al., 2018]. We stress that although in principle our method could be used to investigate the most likely origin of SARS-CoV-2, this would require *Coronaviridae* data of other species (such as pangolins and bats), but current data is biased towards sequences similar to the human SARS-CoV-2 and hence not representative of the original host.

The log-probability difference between the two models is plotted in Fig. 6 as a function of time for the first 1100 days from the start of the 2020 pandemic. It shows a slow but steady adaptation to human nucleotide usage (black line, whose slope is significantly different from 0 with a p-value of 10^*−*9^). Quite surprisingly, the slope of the fitting line is larger for sequences collected in the last 6 months (data downloaded on June 30th, 2023), sug-gesting an increase of the adaptation speed in the Omicron 23A variant that appeared in early January 2023 and rapidly took over the entire SARS-CoV-2 global population. In the above analysis we have taken into account a number of limitations and delicate points that we discuss here. First, the SARS-CoV-2 sequence data is heavily biased, both geographically (a large fraction of the sequences are collected in a small number of countries) and temporarily (the rate of sequence collection increased steadily in the first months of the pandemics). Second, as discussed above, we have used the single Wuhan sequence to infer the model for the unknown virus transmitting host. Third, the time over which the adaptation to the human host has been sampled is much smaller than that of the H1N1 strain, on such a short time scale adaption driven by non-synonymous mutations with clear fitness advantages could result in a confounding signal.

**Figure 6.**
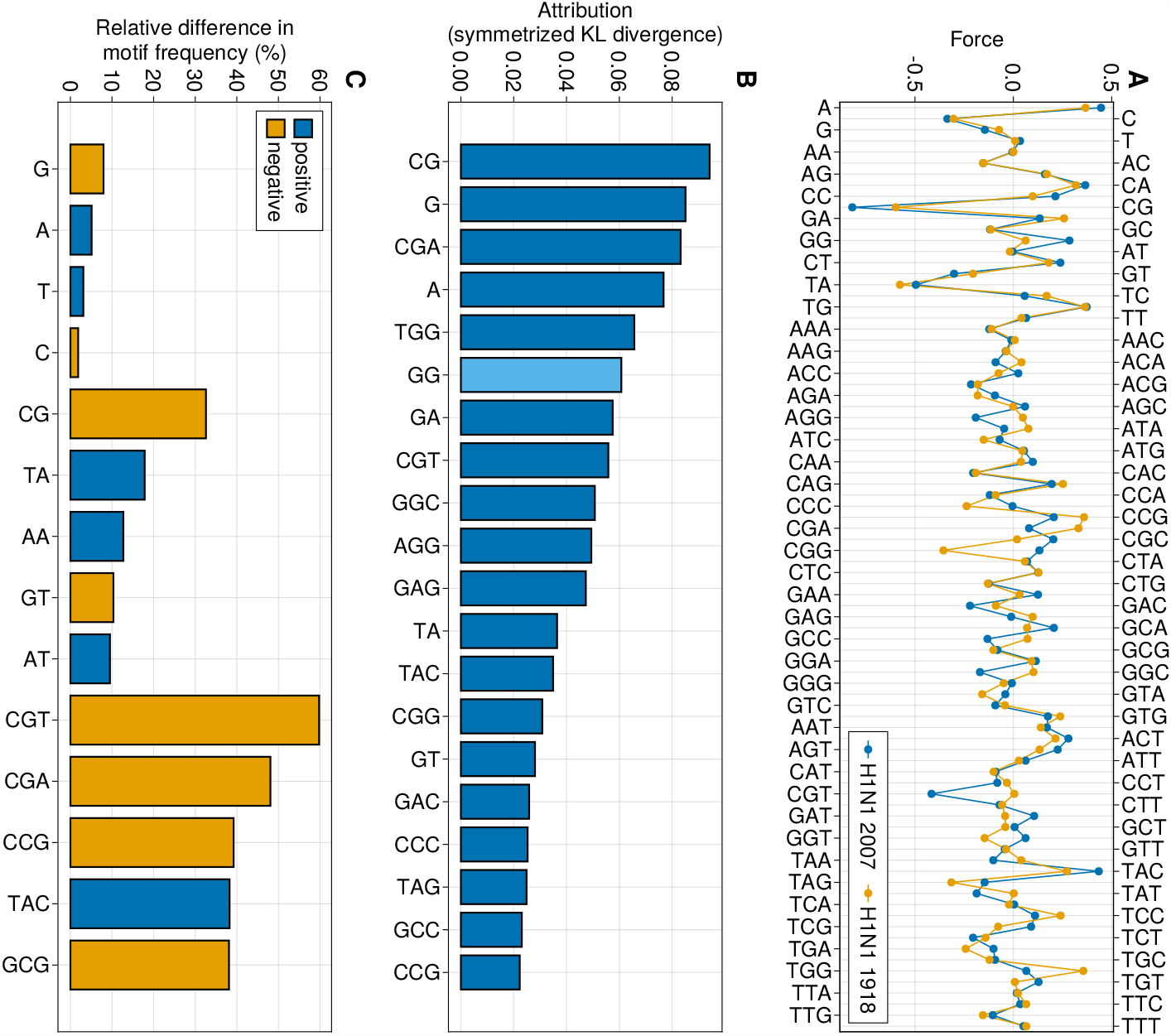
The learned parameters of MENB models can be directly visualized and interpreted. A: Plot of each of the 84 parameters (forces) learned by MENB models trained on all segments but PB2 of H1N1 Influenza A strains collected in 2007 (blue) and of the 1918 strain (orange). B: Attributions computed with the method of integrated gradients (Methods Sec. 5.1.3) for the symmetrized Kullback-Leibler divergence between the MENB models used in panel A. To allow for an easier visualization only the 20 parameters with the highest contribution (in absolute value) to the symmetrized KL divergence are shown. Light blue bars denote negative attributions. C: Relative difference in expected motif frequencies between the MENB models used in panel A (Methods Sec. 5.1.3). Only the 5 top differences (in absolute value) are plotted for 2-mers and 3-mers. Blue (orange) bars correspond to positive (negative) differences.

To address the first issue we used a curated dataset of sequences collected by Nextstrain [Hadfield et al., 2018] to build the model of SARS-CoV-2 in the human host: in this dataset sequences are subsampled to reduce biases from different geographic regions and time periods, and most of the sequences are collected in the last 6 months. As for the second issue, although a MENB model can be trained with a single sequence, in this case the motif frequencies are less representative of the virus-host pair under analysis, so additional caution must be used in this case in interpreting the results obtained. Indeed, by construction the initial log-likelihood associated to the “Wuhan” host will be higher than the one for human *Coronaviridae*. More-over the “Wuhan” host is likely not to be the human [Andersen et al., 2020], but using specific viral sequences that has been collected in bats or pangolins (that have been suggested to be the reservoir of SARS-CoV-2 ancestors) due to their similarity with the Wuhan sequence would give very similar results. Regarding the third problem outlined before, there is no way to deal with it other than collecting sequences for longer times, but the questions of whether some early signals of host adaptation can be spotted with the genomes observed so far is still well-posed.

### 2.5. MENB models’ parameters reflect biologically-relevant features

The MEMB models offer the advantage to have a relatively low number of learnable parameters and that each of them is related to the usage of the corresponding motif. Such models are therefore ideal candidates for interpretation, that in turn can be useful to accumulate insight into potential roles in molecular biology of motifs, for instance associated to the recognition by the host innate immune system.

To showcase this we considered two models trained on the PB2 segments of *Orthomyxoviridae* viruses: one (“H1N1 1918”) has been trained on the sequence collected in 1918, the other (“H1N1 2007”) has been trained on 26 sequences collected in 2007. In Fig. 6A we show the entire parameter profile of the two models. Parameters different form zero reflect the presence of selective forces which push up or down the number of the corresponding motif with respect to sequences generated uniformly at random. Considering the fact that the 1918 strain was likely of avian origin, the first interesting remark is an overall similarly of the force profile in the two cases, especially for nucleotides and dimers, which indicate that many of the force parameters did not significantly change during the adaptation to the human host. The two dinucleotides with the largest negative forces are the CpG, reflecting the well-known avoidance of CpG motifs, followed by UpA, another known avoided motif that is supposed to have a role in codon efficiency [Tulloch et al., 2014, Atkinson et al., 2014]. Moreover, the force in UpG motif is large and positive, likely due to the C*>*U and A*>*G mutational processes on, respectively, CpG and UpA motifs. This observation points out an important concept that is commonly overlooked in k-mer analyses of genetic sequences: the lack of one or more motif is necessarily compensated by an increase in abundance of other motifs, and vice-versa. In our frame-work this is deeply connected to the *gauge choices* that have to be taken due to conservation of probabilities at single, di and tri-nucleotide levels and are discussed in more details in Methods Sec. 5.1.2.

The differences in the parameter profile of In Fig. 6A disclose the selective pressures on the nucleotide biases, dimers and trimers driving the evolution of the viral sequence in the adaptation to the new host. The most striking differences between the 1918 and the 2007 viruses are the further decreases in the CpG force, as well as CGU motifs decrease, from a value around zero in 1918 to a negative value in 2007. An opposite evolution is observed for the GpG force increasing from zero to a positive value and for the CGG force which relaxes from a negative value toward zero in the 2007 (see also Supp. Fig.9). The decrease in *CpG* forces confirms previous findings and what obtained with a simpler model containing only the CpG force, moreover the different behavior for the tri-nucleotide mirrors the context dependence of the CpG loss [Greenbaum et al., 2008, Greenbaum et al., 2014].

A more rigorous way to study the evolution of the forces is to find the key parameters to discriminate the models inferred from the 1918 and 2007 sequences. This problem can be addressed within the framework of integrated gradients [Sundararajan et al., 2017]:We compute the symmetrized KL divergence between the two MENB models as the sum of attributions, i. e. integrated gradients with respect to each parameter (more details about the procedure are given in Methods Sec. 5.1.3; see Suppl. Fig. 7 for the comparison of symmetrized versus non-symmetrized KL divergences). In Fig. 6B we show the values for the top-20 attributions to the symmetryzed KL divergence: consistently with the forces differences, we find that the largest attribution is on CpG dinucleotide, and several 3-nucleotides motifs containng CpG (CGA, CGU, CGG, CCG) are present. The GpG and GpA and UpA dinucleotides and several related trinucleotides (TGG, GGC, GGC, CGG, GAG, TAC, TAG) have a large attribution too.

Once the inference of parameters is performed we can analytically compute the expected number of 1-, 2-, and 3-nucleotide motifs in a viral sequence according to the MENB models (see Methods Sec. 5.1.3), which (as shown in Fig. 5) should reproduce, by model construction, the motif frequencies in the data, as previously shown in Fig. 8A. It is interesting to compare the force attributions in flu evolution to the relative difference in motif frequency Fig. 6C as, due to network effects, they are only marginally related. Nucleotide or dinucleotide usage can, for instance, be driven also by the di-nucleotide and tri-nucleotide forces. In agreement with the force attributions, the CpG dinucletide shows, among all dinucleotides resulting in human-adapted H1N1 strains, the largest relative decrease in 2007 with respect to 1918. Moreover we observe more UA and AA nucleotides with respect to the 1918 strain. As for 3-mers, the signal is dominated by decrease in usage of specific CpG-containing motifs, although for instance an increase of TAC motifs is observed (Fig. 6B). It is important to notice that relative changes of 3-mers cannot be compared immediately with those of 2-mers, due to the fact that there are 64 different 3-mers and 16 2-mers and so individual 3-mers are in general rarer than individual 2-mers and largest changes are to be expected.

We next discuss the force comparison in the context of virus and host classification, from MENB models inferred from the ensemble of sequences for a fixed viral family and host, to bring out similarities and difference in motif usage through the force parameters. The overall similarity of force profiles is again apparent, see Suppl. Fig. 1, reflecting a direct cross contamination and adaptation through zoonotic transmissions or the presence of similar molecular mechanisms driving the adaptation of the viral sequences to the host. Compatibly with Fig. 2 largest differences are present among viruses than among hosts. The attributions and differences in motif usage depends quite strongly on both viral family and pair of host analyzed, as shown in Supp. Fig. 3 and Supp. Fig. 2, further underlying the peculiarities of each viral family and host and the importance of inferring MENB models for each viral family and host independently.

## 3. Discussion

We demonstrate our maximum-entropy approach can successfully be used to predict from a sequence its viral origin and host based on conditional probabilities and Bayes rule. Consistently with some recent empirical observations [Mock et al., 2020], we show viral sequences adapt to the host nucleotide usage under specific viral-family depending constraints. In the host-classification task, our interpretable MENB algorithm has competitive performance with state-of-the-art approaches based on deep neural networks, despite being far simpler in terms of number of learnable parameters. As expected by classical bias-variance trade-off considerations [Posani et al., 2022], our methods is less subject to the specific details of the training data, and shows remarkable out-of-distribution generalization properties. This can be of direct applicability in practical cases, such as when a new viral subfamily is discovered which possesses a genome different enough from those used in the training set. This scenario is likely to become more and more relevant in the near future, as new viral sequences continue to be discovered [Tisza et al., 2020, Edgar et al., 2022].

Our framework can predict the viral genome evolution in a new host, as the log-probability difference in the new host with respect to the previous host increases in time and measures how well the sequence has adapted to its new host environment. This is clearly shown for the H1N1 Influenza for for which we have 100 years of sampled sequences at our disposal; we see a similar trend of host adaptation for the SARS-CoV-2 pandemics as well, which hass accelerated with the expansion of new variants [Di Gioacchino et al., 2021, Kumar et al., 2022].

An important open question is whether the adaptation to the host that we observe directly provides a fitness advantage to the viruses, or if it is a neutral consequence of the viral evolution within a new environment. Arguments for both possibilities exist: for instance, viruses can reduce their CpG content after infection in an host that uses CpG-recognizing antiviral mechanisms (as ZAP in humans) [Shaw et al., 2021], which is likely an adaptation that provides a fitness advantage. On the other hand, the interferon-inducible antiviral protein APOBEC A3G in humans causes hypermutations on cytosines [Chemudupati et al., 2019] and as such it decreases the C content in viral genomes. In this case it is possible that the observed mutations are those that fix in the viral population without destroying the viral life cycle, and so can have null or (extremely) weak replicative fitness effects. The two effects can also coexist and emerge from sequence data on different time scales of the viral evolution. The analysis of the attributions on the early evolution SARS-CoV2 in Supp. Fig (6), shows that among the little changes observed on the overall force parameters, the attributions containing C and U and their repetition (UUU, CCC) are the largest one. These results are cosistent with previous analysis showing the large diminution of *C* occurrences [Hodcroft, 2021] and the presence of local pressures on the CpG, on specific regions of the genome. In particular, large CpG diminution has been observed in the N protein open reading frame which occupies a small region in the genome but is one of the most abundant transcript in the cytoplasm [Di Gioacchino et al., 2021].

The work described here has several potential applications. The fast and flexible host detection algorithm introduced here can easily be integrated within metagenomics studies to infer the host of viruses, even if it is quite different from the sequences used to train the algorithm. Moreover, recent studies have pointed out viral mimicry by some repeats in the human genome, and our group has suggested to use a MENB model to identify similarities between genomic regions and viral families [Šulc et al., 2023]. Secondly, MENB models can be broadly used to study emerging pathogens and their adaptation to new hosts, as a support in surveillance studies. Moreover the modeling at the nucleotide level is necessary to capture some features of viral evolution which should further combined research within the inference of epistatic fitness landscapes of viral genomes that including in a single model synonymous and non-synonymous mutations, as the synonymous mutations may well have fitness costs [Neher and Shraiman, 2011, Zeng et al., 2021]. Finally, thanks to their generative properties underlined here, MENB models are ideal candidate for the optimization in RNA vaccine design for efficiency and minimizing rejection due to immunogenicity [Pardi et al., 2018]. By preditcing how viruses adapt to their new host we can better understand mechanisms that drive their adaptation and design intervention

4 Acknowledgement

This work was supported by grant ANR-19 Decrypted CE30-0021-01, and by the European Union’s Horizon 2020 research and innovation programme under the Marie Sklodowska-Curie grant agreement No 101026293.

## 5. Methods

### 5.1. The maximum entropy nucleotide bias model

In this section, we will first give a maximum-entropy derivation of the MENB model as given in Eq. (2.1). This will clarify why some of the parameters can be arbitrarily fixed as they are redundant (gauge choice) and we will discuss the specific choices in this regard made here. Finally we will describe how all the computations involving the MENB model used in this paper can be performed exactly and efficiently building on classical statistical-physics methods.

#### 5.1.1. Maximum entropy justification

Consider an set of sequences observed (data), we want to find a probability distribution on the sequence space (model) such that: (i) the observed frequencies of nucleotides, 2-mers and 3-mers in the data match those expected by sampling sequences according to the model, and (ii) the entropy ∑_*s*_ *p*(*s*) log *p*(*s*) is maximized. Therefore the MENB model probability distribution maximizes the following quantity

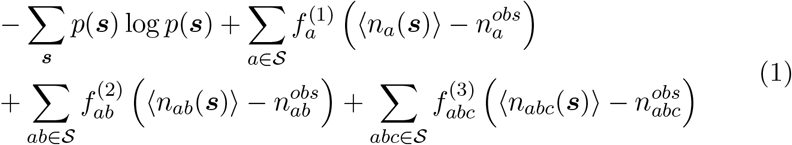

over *p*(***s***) and the Lagrange multipliers 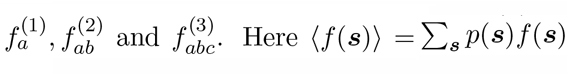, and quantities with the *obs* superscript are averages computed on the data sequences. By taking the functional derivative with respect *p*(***s***), we obtain the functional form given in Eq. (2.1), where the Lagrange multipliers, that we also call force parameters, need to be fixed so that the observed frequencies of nucleotides, 2-mers and 3-mers in the data match those expected by sampling sequences according to the model. Follwing [Greenbaum et al., 2014], this parameter inference can be performed by computing the partition function

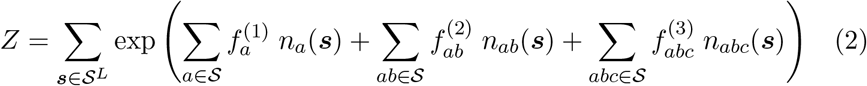

that normalizes the probability distribution in Eq. (2.1) and using it to estimate the quantities ⟨*n*_*a*_(***s***)⟩, ⟨*n*_*ab*_(***s***)⟩, ⟨*n*_*abc*_(***s***)⟩. Finally, a root-finding algorithm such as the Newton–Raphson method can be used to find the correct values for the parameters. Optionally the observed quantities 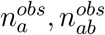 and 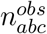 can be regularized by adding pseudocounts to avoid parameter divergences or to give less weight to the sequence details during the inference.

#### 5.2.1. Gauge choices for MENB model

The MENB model specifies a probability distribution over sequences of length *L*. As such, any change of parameters that does not change the probability of any sequence does not have any observable effect and it is called a gauge degree of freedom. For instance, we can send 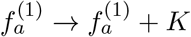 and, for any value of *K*, this modification does not impact the probability of any sequence as it can be readily showed using the fact that 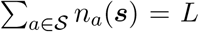. As a consequence, we are free to choose a value for *K* so that, for instance, 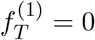, or so that 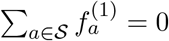

The presence of gauge degrees of freedom stems from the fact that there are many ways of choosing the 84 force parameters in Eq. (2.1) so that the observed frequencies of nucleotides, 2-mers and 3-mers in the data match those expected from to the model. Indeed, although this requirement can be written as a set of 84 equations, some of them are not independent because of the following considerations:

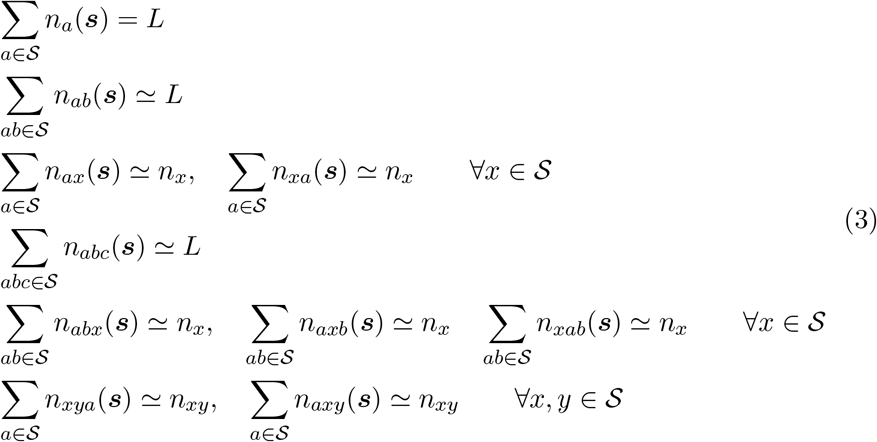

where the symbol ≃ means that the condition is respected in the large-*L* limit, which is the relevant case for all sequences considered in this work. This set of equations can be used to fix the gauge degrees of freedom (“choose the gauge”), and we do so in this work by choosing a gauge where the maximum number of parameters is set to zero, that we call lattice-gas gauge (with a slight abuse of notation), or by choosing a gauge where there is no arbitrary symmetry breaking among the model parameters, that we call zero-sum gauge.

For the lattice-gas gauge, we decide to set to zero all forces of the form 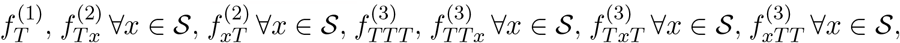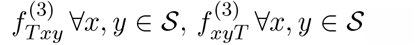. Therefore non-zero *T* -containing forces only have the form *h*_*xT y*_ with *x, y* ∈ S. This means that the effective number of free parameters to be inferred goes from 84 to 48.

The lattice-gas gauge is particularly useful to speed-up the inference process and to avoid the Newton–Raphson method to fail to converge due to flat directions in the parameter space, but it is not practical when looking at the inferred parameters to interpret them. For this reason after inference we use the zero-sum gauge, that is defined by the following set of equations

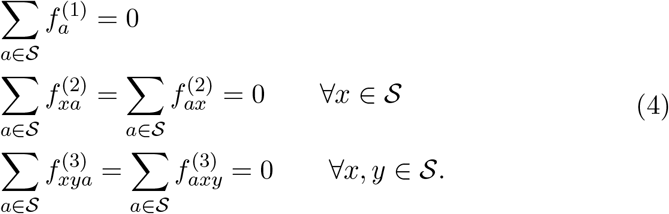

#### 5.1.3. Computation of the partition function and related quantities

An remarkable characteristic of the MENB model is that the partition function *Z* given in Eq. (2) can be computed exactly in a time that scales linearly with the length of the sequence *L* using the so-called transfer matrix method, well-known in statistical physics. This method has been already described for a similar problem in [Greenbaum et al., 2014] (Supporting Information), and the only difference in this case is that the matrices also contain a term that accounts for the 3-body interaction.

Once the partition function of a MENB model is computed, we have immediate access to a wealth of relevant quantities. In particular, we can compute the expected number of *ℓ*-mers *M* as

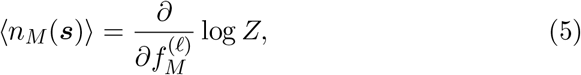

which is the main quantity used to produce Fig. 6B.

Another relevant quantity is the Kullback-Leibler divergence between two models, *p*_1_ and *p*_2_. It can be written as

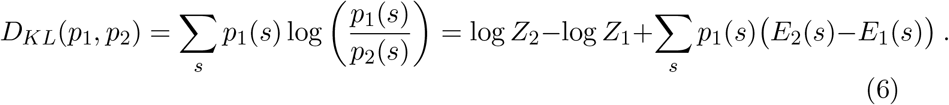

log *Z*_1_ and log *Z*_2_ can be computed exactly with the transfer matrix method, and to compute the last term on the r.h.s. of Eq. (6) we define

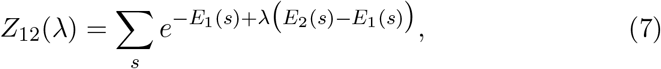

and we have

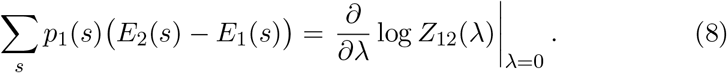

From the KL divergence we can compute the attributions showed in Fig. 6C. Following [Sundararajan et al., 2017], we consider two MENB models defined by the force parameters ***f***_1_ and ***f***_2_. We will use the notation *D*_*KL*_(***f***_1_, ***f***_2_) to denote the KL divergence between the models with parameter ***f***_1_ and ***f***_2_. Thanks to the fundamental theorem of calculus for line integrals, and using *D*_*KL*_(***f***_1_, ***f***_1_) = 0, we get

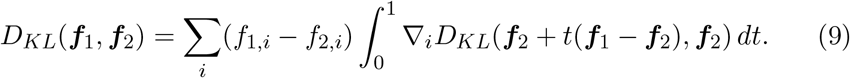

The individual terms of the sum in this equations are the attribution plotted, after rescaling for the total KL divergence, in Fig. 6C and Suppl. Fig. 3. As a final remark, we notice that the attributions depends on the gauge used. In this work we always computed attributions in the zero-sum gauge, and we observe that if the parameters ***f***_1_ and ***f***_2_ are from models in the zero-sum gauge, then Eqs. (4) still hold for ***f***_1_ + *t*(***f***_1_ − ***f***_2_), and so the path of models used in Eq. (9) preserve the zero-sum gauge.

### 5.2. Data and code availability

All sequence data has been collected from the BV-BRC database [Olson et al., 2022]. After discarding short viral sequences (length lower than 1000 bases), we selected the pairs of host and viral family so that each viral family has at least 100 sequences annotated with each host chosen. We discarded Influenza A sequences collected after 2009 as the database is dominated by strains of the H1N1 “swine flu”, whose triple-reassortment origin [Garten et al., 2009] and (likely) not perfect adaptation to humans is a confounding factor during training. The resulting dataset, that is the starting point for all the results presented here, is available at https://zenodo.org/doi/10.5281/zenodo.10050076. The SARS-CoV-2 data used for Fig. 5B can be downloaded at https://nextstrain.org. Notice, however, that this data is often updated as it is focused on the last 6 months. To allow exact reproducibility of our results we uploaded the data we used (downloaded on June 30th 2023) together with the sequence data at https://zenodo.org/doi/10.5281/zenodo.10050076.

The code to infer models is written in Julia and publicly available in the GitHub repository at https://github.com/adigioacchino/MaxEntNucleotideBiases.jl.

We trained our MENB models on 100 viral sequences randomly selected from our dataset for each pair of host and viral family. We replicated this sub-sample three times, observing quite small quantitative differences based on the sequence choice (see error bars in Fig. 1). For the comparison with VIDHOP presented in Fig. 1A, C we used exactly the same sequences. A Snakemake pipeline to train and test the MENB models on the data used here is available at https://github.com/adigioacchino/MENB_snakemake.

## 6. Supplementary figures

**Suppl. Fig 1:**
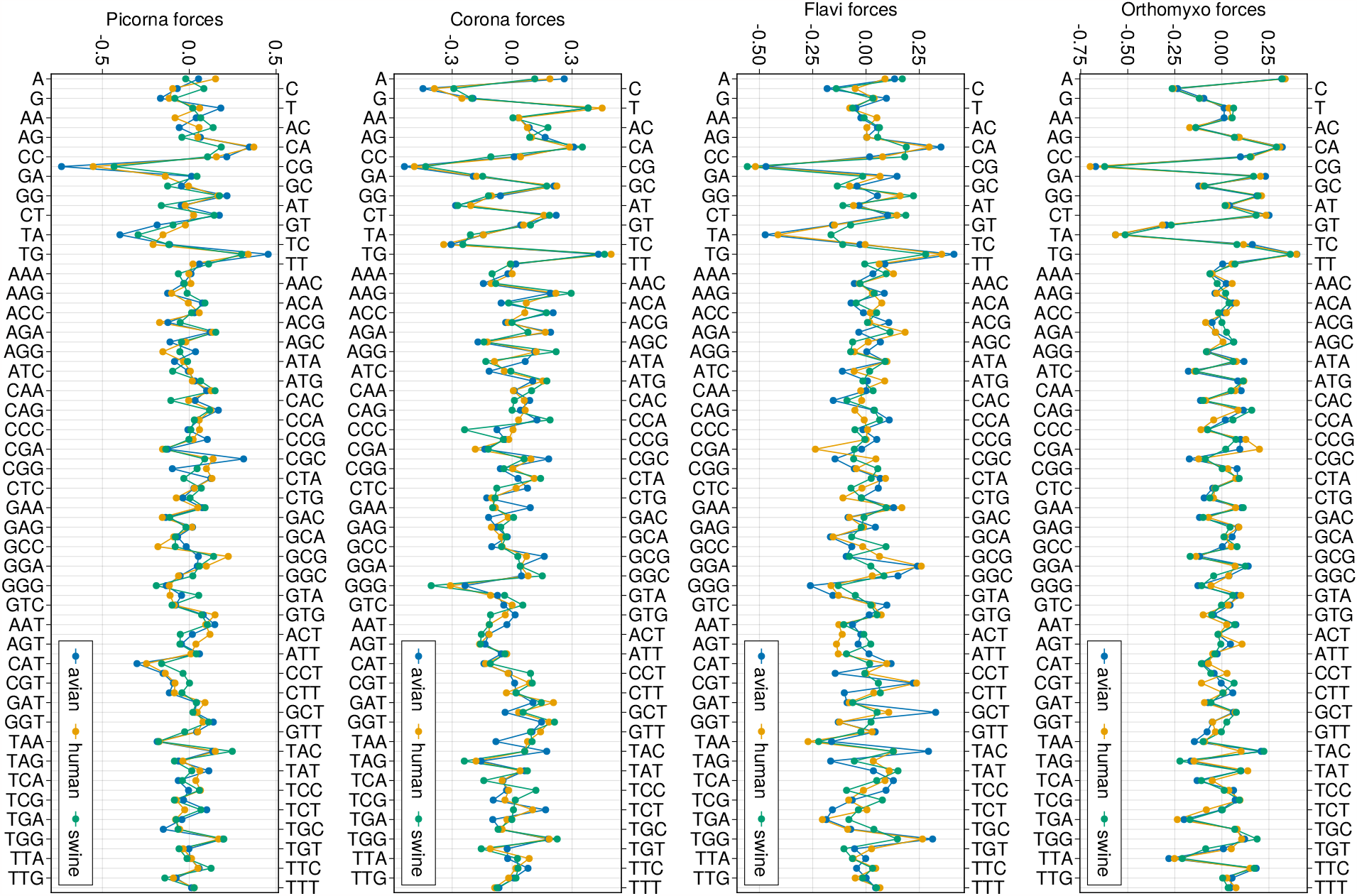
All forces shown for each model learned in this work.

**Suppl. Fig 2:**
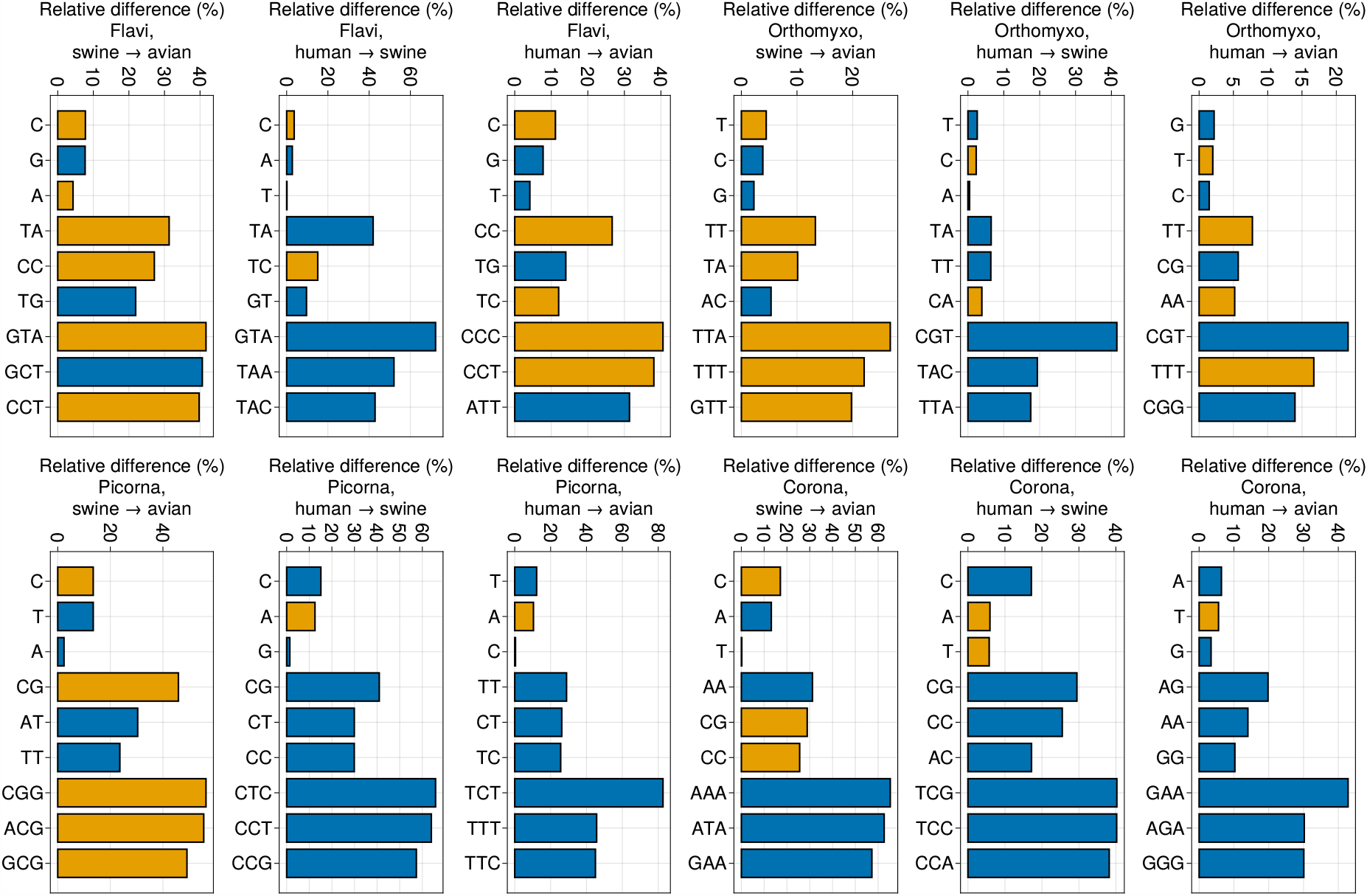
Relative difference in motif usage shown for each pair of hosts at given viral family. Blue bars correspond to increases in motif usage, and orange bars to decreases. Only the 3 highest differences (in absolute value) are shown for nucleotides, 2-mers and 3-mers.

**Suppl. Fig 3:**
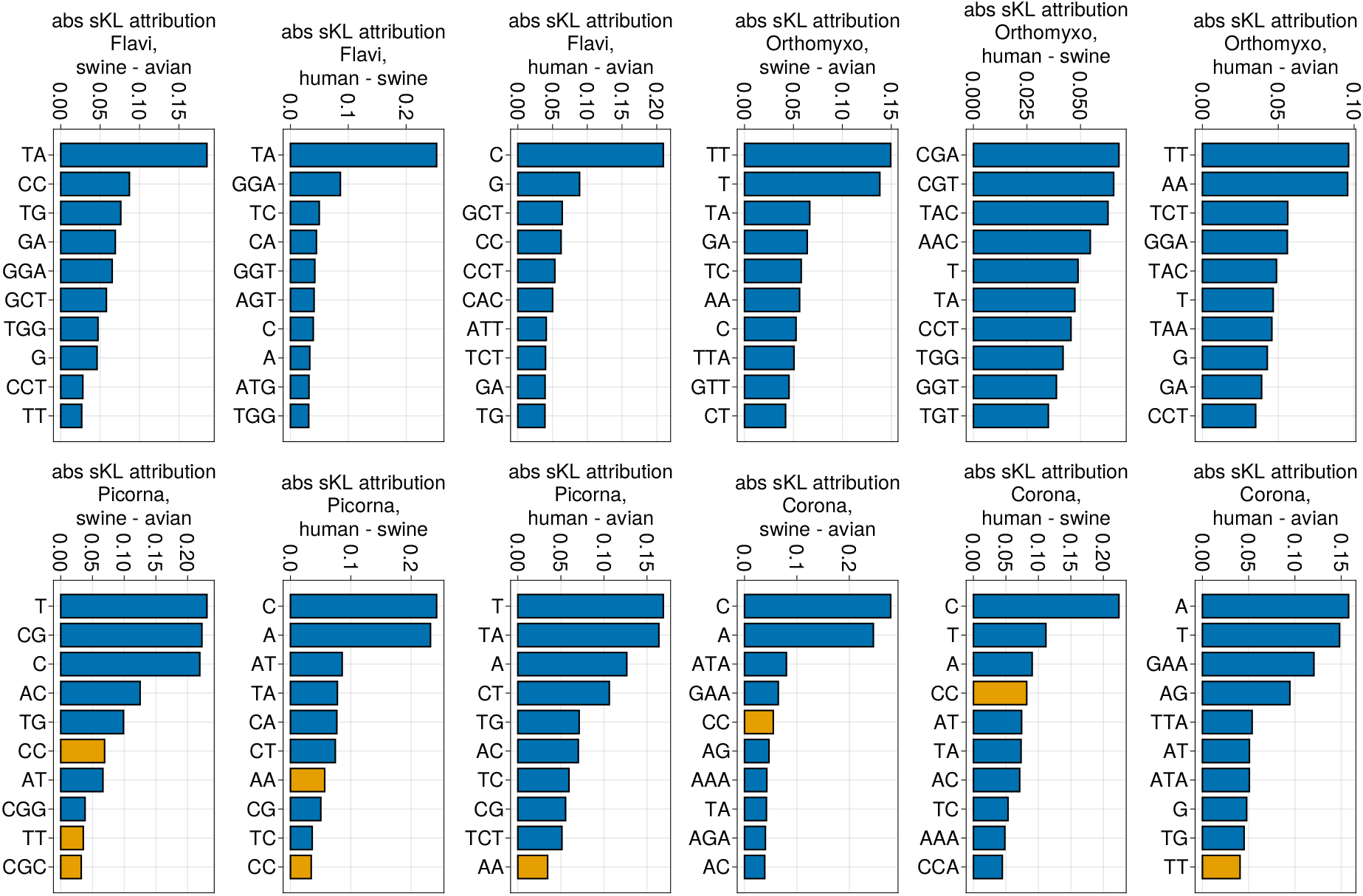
Attribution to symmetrized KL divergence shown for each pair of hosts at given viral family. Blue bars correspond to positive attributions, and orange bars to negative attributions. Only the 10 highest attributions (in absolute value) are shown.

**Suppl. Fig 4:**
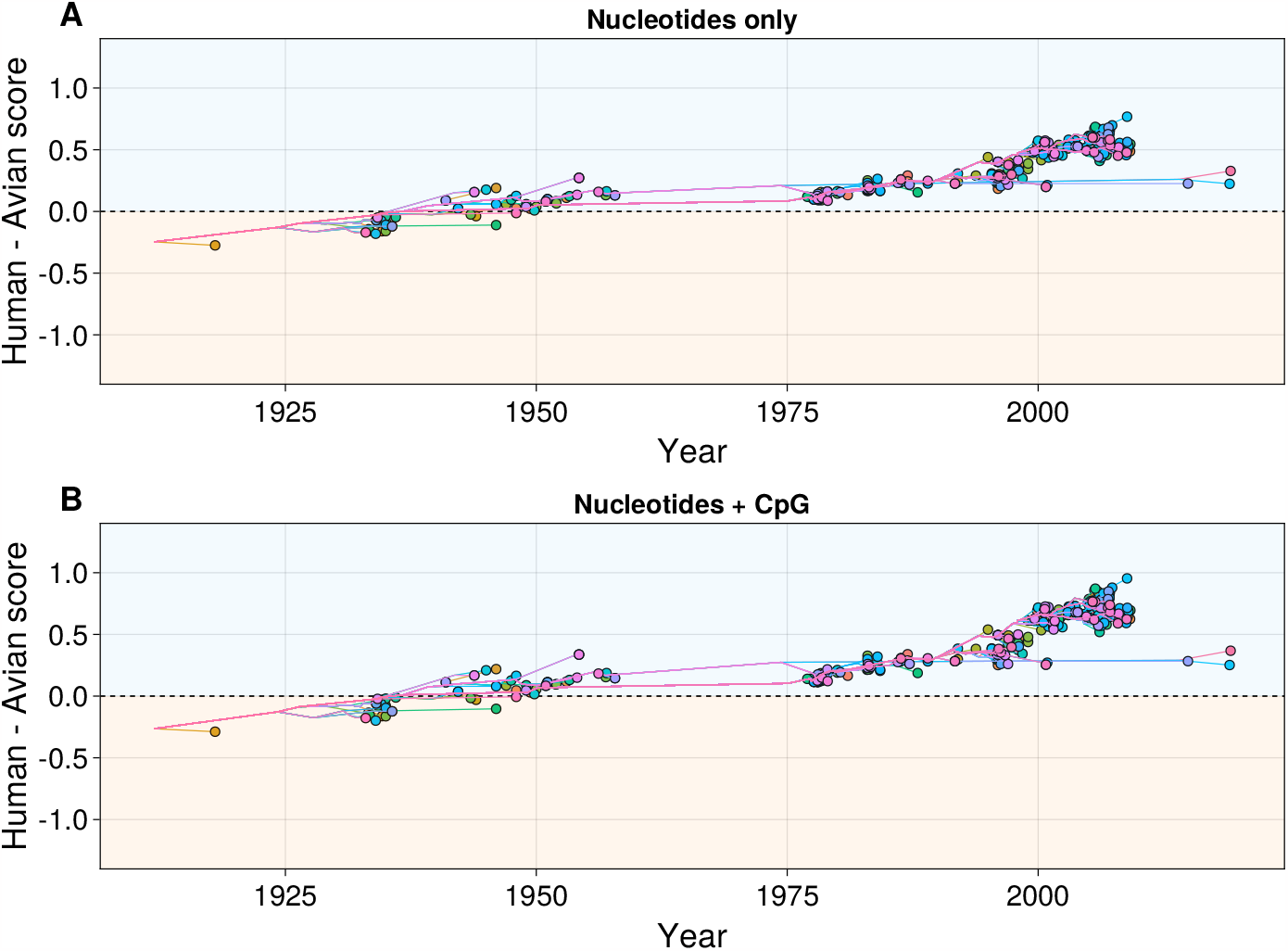
Loglikelihood differences of simplified MENB *Orthomyxoviridae*humand and avian models versus time of H1N1 Influenza A sequences. In panel A a model with only nucletide force inferred is used, and in panel B these forces are inferred together with the CpG force. The colored lines are the reconstructed paths of the inferred phylogenetic tree that connect the root to each leaf (observed sequence), and the score versus inferred time is plotted also for the internal node (inferred) sequences.

**Suppl. Fig 5:**
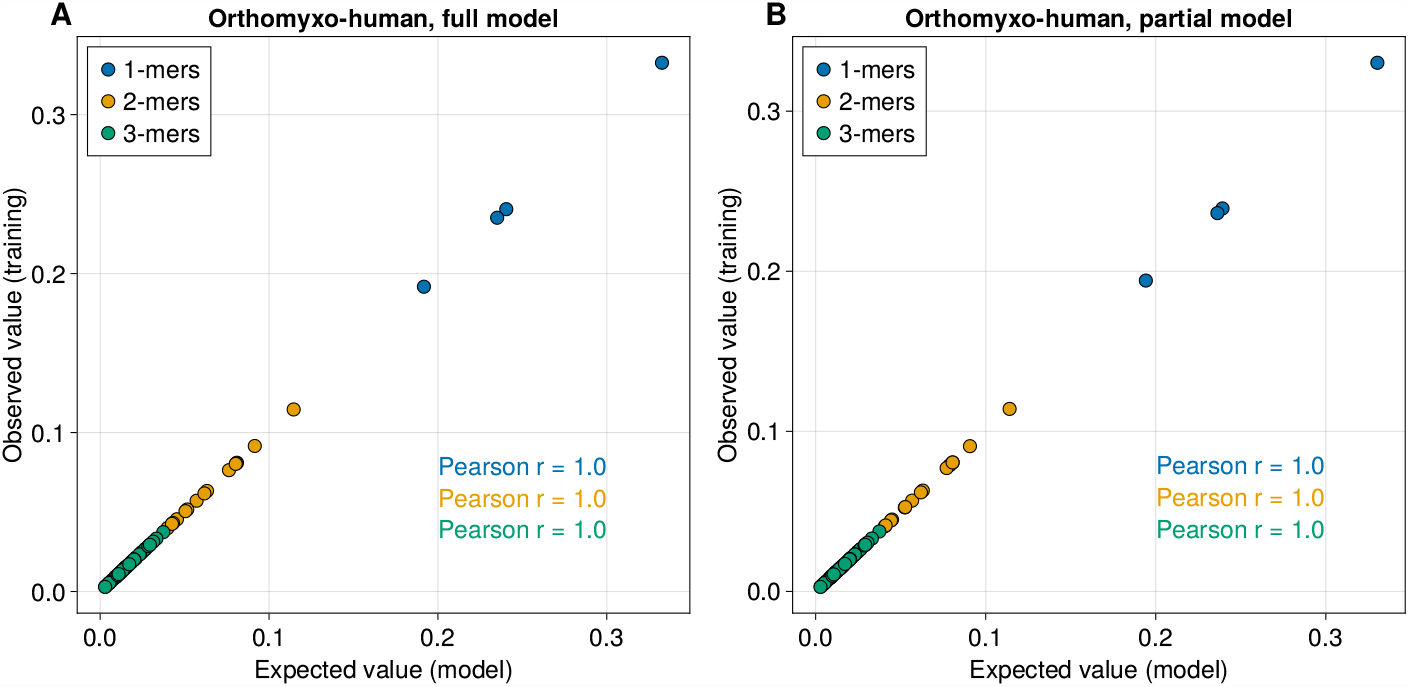
A: Frequency of nucleotides, 2-mers and 3-mers observed in the training set of full human *Orthomyxoviridae*sequences versus the value obtained analytically from the inferred MENB model. B: Same as A for the MENB model trained on human *Orthomyxoviridae*sequences without the segment coding for PB2. .

**Suppl. Fig 6:**
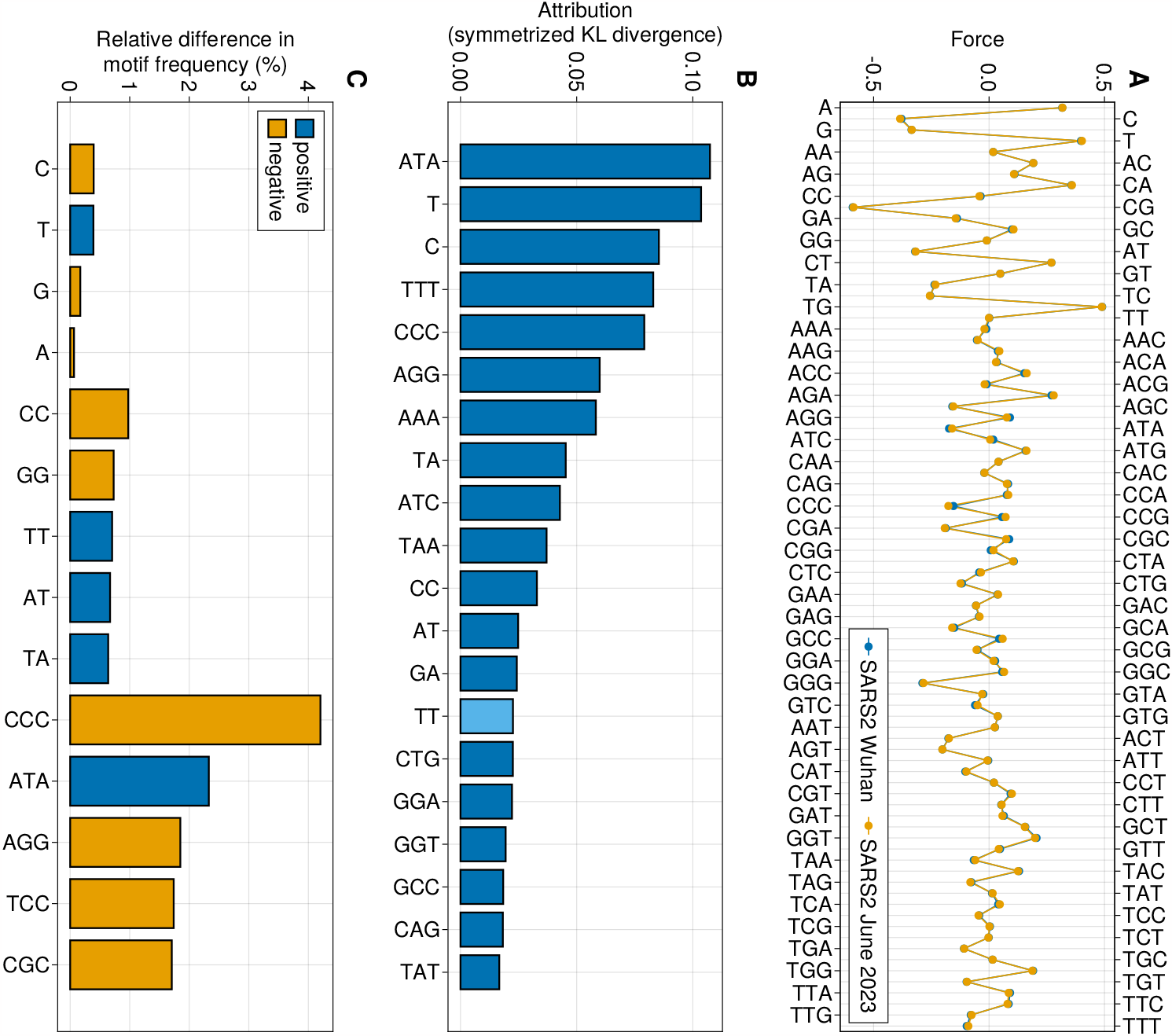
A: Plot of each of the 84 parameters (forces) learned by MENB models trained on the SARS-CoV-2 sequence collected in Wuhan in December 2019 (blue) and on sequences collected in June 2023 (orange). B: Relative difference in expected motif frequencies between the MENB models used in panel A (Methods Sec. 5.1.3). Only the 5 top differences (in absolute value) are plotted for 2-mers and 3-mers. Blue (orange) bars correspond to positive (negative) differences. C: Attributions computed with the method of integrated gradients (Methods Sec. 5.1.3) for the symmetrized Kullback-Leibler divergence between the MENB models used in panel A. To allow for an easier visualization only the 20 parameters with the highest contribution (in absolute value) to the symmetrized KL divergence are shown. Orange bars denote negative attributions.

**Suppl. Fig 7:**
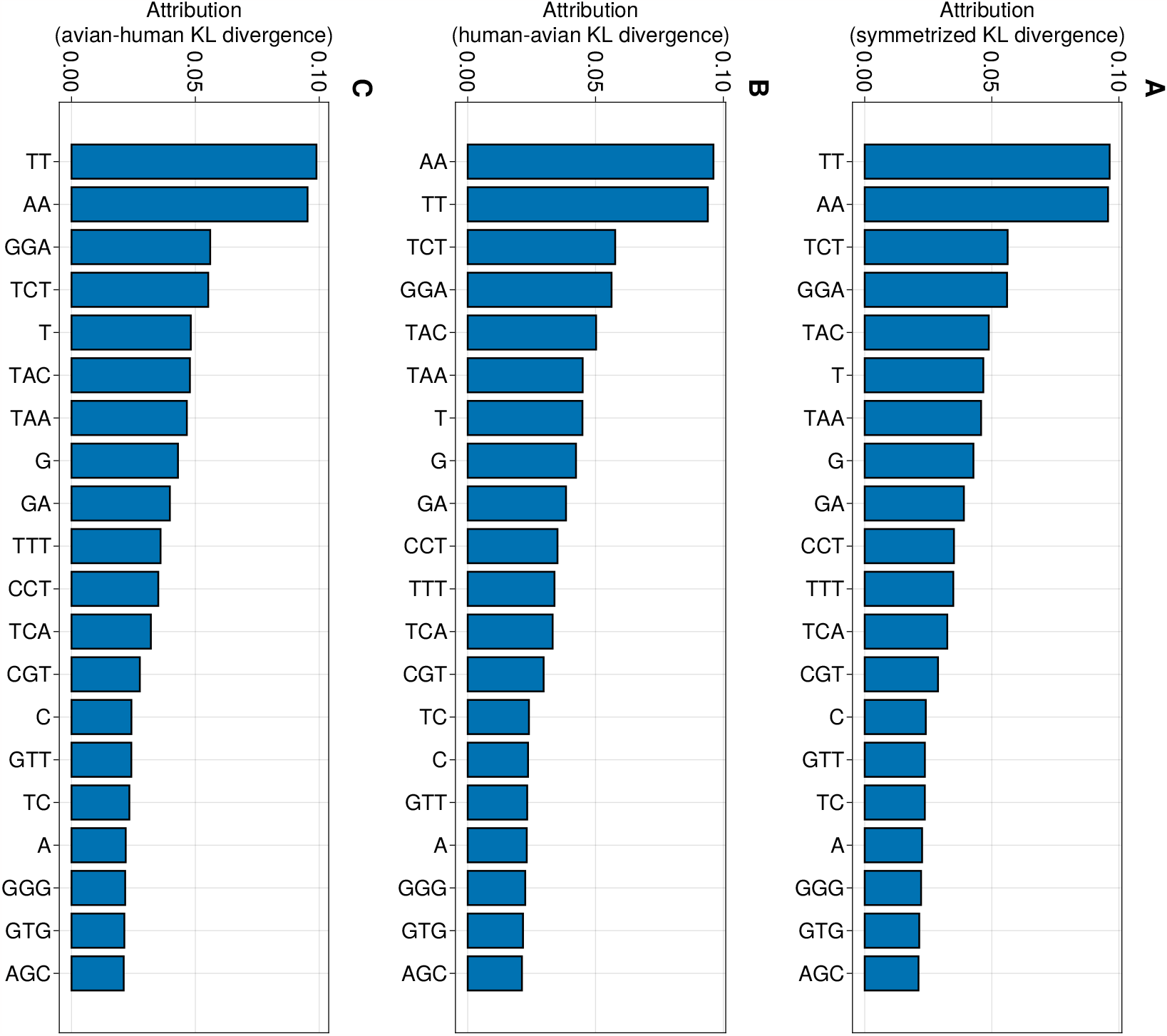
Comparison between attribution to the symmetrized KL divergence between *Orthomyxoviridae* human and avian viruses (panel A), and the two non-symmetrized KL divergences that compose it (panels B, C).

**Suppl. Fig 8:**
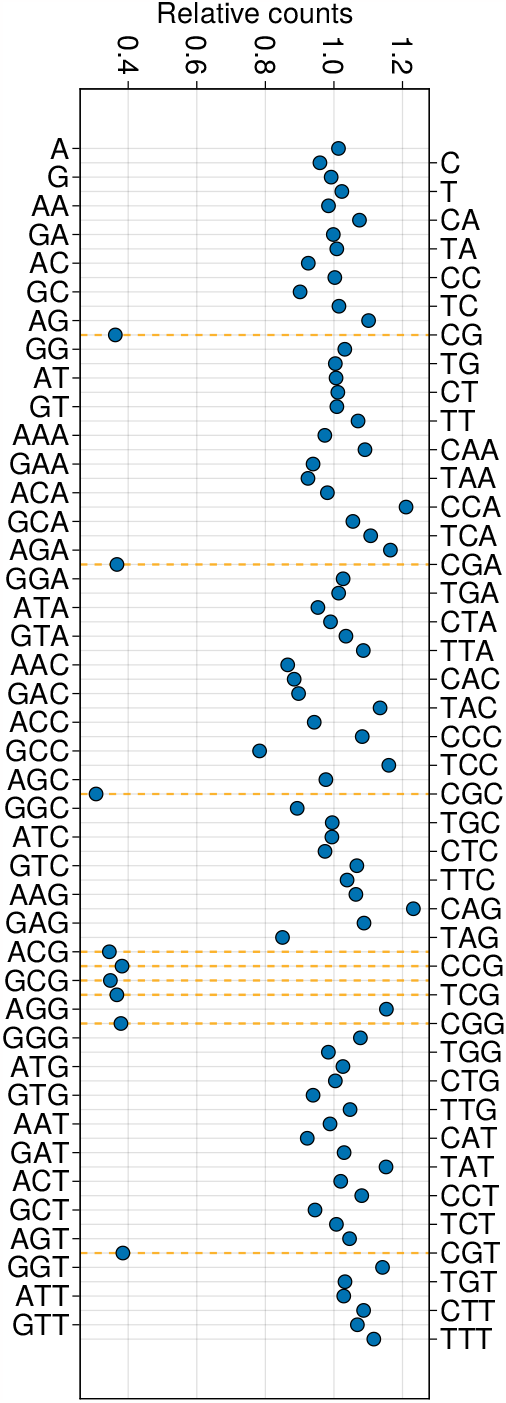
Comparison between the number of motif observed in the 1918 H1N1 PB2 sequence and in PB2-coding sequence synthetically evolved to reduce their CpG number. A value of 1 means no change in motif abundance. CpG-containig motifs are highlighted with orange lines.

**Suppl. Fig 9:**
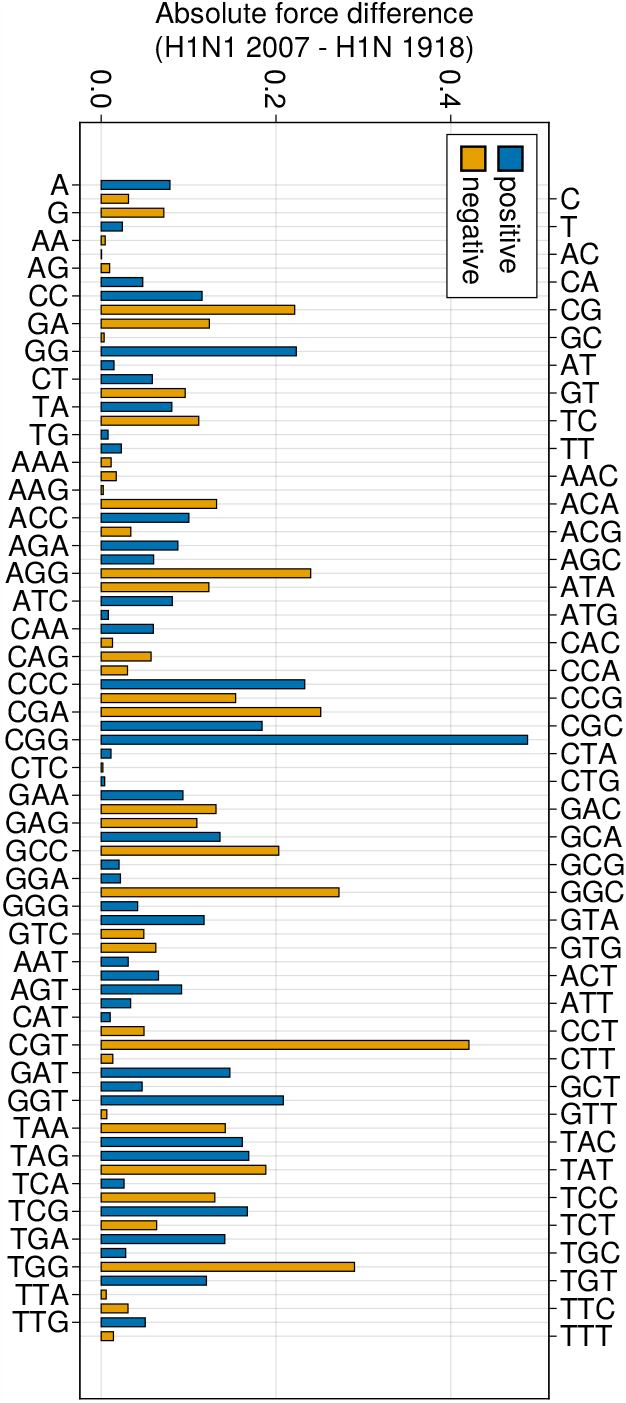
Comparison between the forces inferred on the 1918 and in 2007 H1N1 sequences. Blue/orange bars correspond to increased/decreased forces of 2007 sequences with respect to the 1918 sequence.

